# Human PAR1 expressed on mouse platelets contributes to hemostasis and arterial occlusion

**DOI:** 10.1101/2025.06.03.657740

**Authors:** Sachiko Kanaji, Alessandro Zarpellon, Roberto Aiolfi, Yosuke Morodomi, Jennifer Orje, Antonella Zampolli, Eric Won, Zaverio M. Ruggeri, Taisuke Kanaji

## Abstract

Thrombin (FIIa) signaling through protease-activated receptors (PARs) is a relevant platelet activation mechanism. Human (h) PAR1 antagonists are approved for antithrombotic therapy, but bleeding is a concerning complication. In addition to PARs, platelet glycoprotein (GP) Ibα also binds FIIa enhancing human platelet response to lower agonist concentrations *ex vivo*. Signaling through GPCRs is well understood, but how GPIbα association with distinct PARs modulates FIIa-dependent platelet activation *in vivo* remains unclear. One obstacle in addressing this question is the distinct human platelet PAR1/PAR4 expression as opposed to mouse (m) PAR3/PAR4. Previous attempts to express functioning hPAR1 in mouse platelets using platelet-specific promoters or targeting into the mPAR3 locus have not been successful. Here we report our studies using a hPAR1 transgene with a floxed STOP sequence and strong synthetic (CAG) promoter targeted into the mouse Rosa26 locus. Generated Rosa26-hPAR1Tg^fl^ mice were then sequentially crossbred with PF4–Cre and PAR3^−/−^ mice, yielding mP3^−/−^hP1Tg mice whose platelets expressed hPAR1 with endogenous mPAR4. These mice, unlike PAR3^−/−^, had no excessive bleeding after tail clipping, but bled profusely after administration of the hPAR1 antagonist, vorapaxar. Accordingly, mP3^−/−^hP1Tg mice had more frequent and stable post injury occlusion of the carotid artery than PAR3^−/−^ mice, but this difference was abolished by vorapaxar treatment. We anticipate that studies in this mouse strain will help unravel the regulation of PAR-mediated thrombin–induced platelet activation *in vivo* with findings more directly relevant to human pathophysiology.

**Key Points (<140 characters each):**

- Human (h) PAR1 expressed in the platelets of PAR3^−/−^ mice shortens the bleeding time and promotes more frequent and stable post injury carotid artery occlusion.
- The specific hPAR1 antagonist drug, vorapaxar reverses the phenotypic changes associated with hPAR1 expression in PAR3^−/−^ mice expressing mouse PAR4.
- Experimental data from PAR3^−/−^ mice expressing hPAR1 should be directly relevant to the development of specific thrombin-induced platelet activation inhibitors.

## Introduction

Thrombin (FIIa) is a multifunctional serine protease that cleaves fibrinogen, leading to fibrin clot formation, and evokes multiple effects on various cell types, including platelets and other vascular cells. FIIa cellular effects are mediated by protease-activated receptors (PARs), members of the family of G-protein-coupled receptors (GPCR).^1^ Human platelets express PAR1 and PAR4, which couple to the same set of G proteins and signaling molecules but exhibit functional differences. The first extracellular PAR1 domain contains a hirudin-like sequence, absent in PAR4, to which FIIa binds with high affinity inducing rapid platelet responses with higher sensitivity compared to PAR4.^2^ The latter is activated by higher FIIa concentrations and induces slow and prolonged signaling to sustain irreversible aggregation.^3,4^ Mouse platelets, in contrast, express PAR3 and PAR4, of which PAR3 does not transduce signal but acts as a co-receptor enhancing PAR4 response to FIIa stimulation.^5^ The species-specific difference in platelet PAR expression has hampered the use of mouse models to study physiopathologic significance and pharmacologic targeting of the FIIa-PAR pathway of platelet activation in hemostasis and thrombosis.

In addition to PARs, the GPIbα amino-terminal domain on human platelets contains high-affinity FIIa binding sites and is thought to contribute to platelet activation^6,7^ and platelet procoagulant activity.^8^ X-ray crystallography has revealed that binding to GPIb*α* involves both FIIa exosite I and II,^9-11^ such that one FIIa interacts with two GPIbα molecules and each GPIbα, in turn, interacts with two FIIa molecules.^12^ FIIa-GPIbα binding supports activation of human platelets via PAR1, but mechanisms and functional significance of this cooperation are yet to be fully explained.^13-15^ In addition, it is still controversial whether FIIa-GPIbα interaction enhances or inhibits platelet activation via PAR4.^12,16,17^ Thus, we set out to generate a mouse model with human-like PAR1/PAR4 expression and function as a tool to clarify unresolved issues relevant to FIIa-mediated platelet activation.

## Methods

### Animal experiments

C57BL/6J and PF4-Cre mice were from The Jackson Laboratory. Par3 knockout mice (P3^−/−^) were a kind gift from Dr. Shaun Coughlin, University of California at San Francisco.

Complete blood counts (CBC) were obtained using a Procyte Dx instrument (IDEXX Laboratories, Westbrook, ME). All experimental animal protocols were approved by the Scripps Research Institutional Animal Care and Use Committee (IACUC).

### Human subjects

Blood samples were obtained in accordance with the Declaration of Helsinki from healthy donors who signed an informed consent. All experimental human protocols were approved by The Scripps Research Institute Institutional Review Board.

### Generation of transgenic mice expressing human PAR1

A targeting vector was constructed by cloning the hPAR1 cDNA into the CAG-STOP-eGFP-ROSA26TV plasmid (CTV, Addgene, plasmid#15912).^18^ This was electroporated into mouse ES cells and correctly targeted clones were used for blastocyst injection, resulting in chimeric mice that were bred with C57BL/6J females to confirm germline transmission. Transgenic (Rosa26-hPAR1Tg^fl^) and PF4-Cre mice were crossed to remove the STOP sequence and induce expression in megakaryocytes (MKs) and platelets. Both Rosa26-hPAR1Tg^fl^ and PF4-Cre strains were bred into P3^−/−^ background to generate P3^−/−^hPAR1Tg mice. The same strategy, but with a targeting hGPIbα vector in the same CTV plasmid, was used to generate hGPIbα mice on a mouse GPIbα^−/−^ background for methodological control and future studies.

### Platelet functional assays

Mouse blood was collected by retro-orbital bleeding into 1/10^th^ volume of 0.113 M sodium citrate as anticoagulant, then diluted 1:1 with Tyrode’s buffer (pH 6.0) before centrifuging at 230 g for 7 minutes. The resulting platelet-rich plasma (PRP) was separated from sedimented red cells and transferred into a tube containing 3 times the blood volume of Tyrode’s buffer with 0.0113 M sodium citrate (pH 6.0). After centrifugation at 1,000 g for 12 minutes, the resulting platelet pellet was suspended into Tyrode’s buffer (pH 7.4) at a platelet count adjusted to 300·10^3^/μL. For studies with human samples, platelets from healthy donors washed as detailed above were prepared at 250·10^3^/μL. Platelet aggregation was measured with a Model 440 aggregometer (Chrono-Log Corporation). For intracytoplasmic Ca^2+^ concentration ([Ca2^+^]_i_) assays, washed platelets loaded with 5 μM Fura 2-AM were stimulated with FIIa in the presence or not of specified inhibitory antibodies and fluorescence intensity was recorded at 510 nm with alternating excitation at 340/380; the ratio of the respective values was converted to Ca^2+^ concentration.^19,20^

### Western blot analysis

Washed platelets were resuspended in Tyrode’s buffer and lysed with an equal volume of T-SDS buffer (50 mM Tris-Cl [pH 7.4], 2% SDS) in the presence or absence of β-mercaptoethanol. The lysates were subjected to SDS-PAGE using either 7.5% or 4–20% polyacrylamide gels. Proteins were then transferred electrophoretically to polyvinylidene fluoride (PVDF) membranes (Millipore). Membranes were blocked with TBS-T (Tris buffered saline with Tween 20) containing 5% BSA and incubated with primary antibodies, followed by IRDye 680 or 800-conjugated goat anti-mouse or goat anti-rabbit IgG secondary antibodies. Protein bands were visualized using the LI-COR Odyssey imaging system, and band intensities were quantified using Image Studio software (LI-COR Biosciences).

### Flow cytometry assays

Mouse blood samples were stained with Brilliant Violet 421™ labeled anti-CD41 antibody (MWReg30, BioLegend, San Diego, CA) to gate the platelet population and Alexa Fluor 647 labeled anti-human PAR1 antibody (ATAP2, Santa Cruz Biotech, Santa Cruz, CA). For MK analysis, mouse bone marrow (BM) cells were fixed in 1% paraformaldehyde at 4°C for 2 hours. Cells were then washed, resuspended in phosphate-buffered saline (PBS, pH 7.4) and incubated for 10 minutes at room temperature (22-25 °C) with 1 µg/mL Brilliant Violet 421™ anti-CD41 antibody (BioLegend) and 100 µg/mL propidium iodide (Sigma-Aldrich, MO). Samples were analyzed using a Novocyte flow cytometer (ACEA Biosciences, CA) and the results were evaluated using FlowJo software (FlowJo, LLC).

### Tail bleeding time assay

Mice were anesthetized with isoflurane administered in a precision vaporizer before clipping 3 mm of the tail distal tip with a sterile scalpel blade. The injured tail was immersed into isotonic saline at 37° C and the time to cessation of blood flow was recorded (bleeding time). Hemorrhage persisting after 600 seconds was stopped by cauterizing the tail wound.^21^

### In vivo arterial occlusion

The common carotid artery of anesthetized mice was dissected, and an ultrasound flow probe was positioned around the vessel.^22^ After measuring baseline flow, a silicon strip is positioned underneath the artery and a 0.7-μL drop of 0.22 M FeCl_3_ solution was applied to the surface of the adventitia for 3 minutes. After removing the silicone strip and washing the surrounding area, the carotid blood flow was monitored for the total observation period of 33 minutes. The artery was considered occluded when flow was <0.1 mL/min. Stable occlusion was defined as flow rate < 0.1 mL/min for at least 10 minutes. A flow index was calculated as the ratio between the volume of blood that flowed through the carotid artery during the observation time and the volume calculated, assuming a constant flow rate equal to the maximum value observed during the first minute after injury.

### Statistical analysis

Results were analyzed using Prism (GraphPad Software, La Jolla, CA; version 9.3.1 and higher) as detailed in the figure legends.

## Results

### Distinct effects of FIIa-GPIbα binding on PAR1 and PAR4 signaling in platelets

In normal human platelets, the anti-GPIbα monoclonal antibody (MoAb), LJ-Ib10, and the function-blocking anti-PAR1 MoAb, WEDE, inhibited the transient [Ca^2+^]_i_ increase indicative of activation induced by FIIa at low concentration (0.5 nM; Figure 1A, left panel) in similar manner. The inhibitory effect of LJ-Ib10 was lesser at intermediate (2 nM) and absent at higher (10 nM) FIIa concentrations, while WEDE delayed [Ca^2+^]_i_ increase even in response to 10 nM FIIa (Figure 1A, middle and right panel). In contrast to inhibiting, LJ-Ib10 enhanced the response elicited by low-dose (0.25 nM) FIIa in platelets from transgenic mice with human (h) GPIbα replacing the wild type homologue (hGPIbα-WT strain;^22^ Figure 1B, left panel). The enhancement was less apparent with 2 nM and disappeared with 10 nM FIIa stimulation as the response of control platelets increased (Figure 1B, middle and right panels). It was previously shown that mouse platelets expressing human GPIbα mutated to lack FIIa binding (hGPIbα-D277N) had enhanced FIIa-dependent *ex vivo* and *in vivo* PAR4-mediated platelet responses compared to hGPIbα-WT mice^17^. Accordingly, hGPIbα-D277N platelets stimulated by 0.25 nM FIIa had enhanced [Ca^2+^]_i_ increase compared to hGPIbα-WT platelets, and the response to low and increasing FIIa concentrations almost exactly overlapped that of hGPIbα-WT mouse platelets treated with MoAb LJ-Ib10 (Figure 1B). Altogether, these findings indicated that FIIa-hGPIbα binding enhances PAR1 signaling in human platelets but decreases PAR4 signaling in mouse platelets. Thus, we reasoned that a “humanized” mouse model with platelets expressing PAR1/PAR4 rather than PAR3/PAR4 could greatly help understand how GPIbα regulates FIIa-dependent platelet functions relevant to human pathobiology. To this end, we targeted hPAR1 cDNA to the mouse *Rosa26* locus, a safe-harbor for transgene expression.^23^ The targeting construct (Figure 1C) contained the cytomegalovirus immediate enhancer/β-actin (CAG) promoter to drive high expression;^24^ a floxed STOP cassette to allow tissue-specific expression upon excision by Cre recombinase;^18,33^ and Internal Ribosome Entry Site (IRES) sequences, enabling EGFP co-expression to facilitate offspring screening.^25^

**Figure 1.**
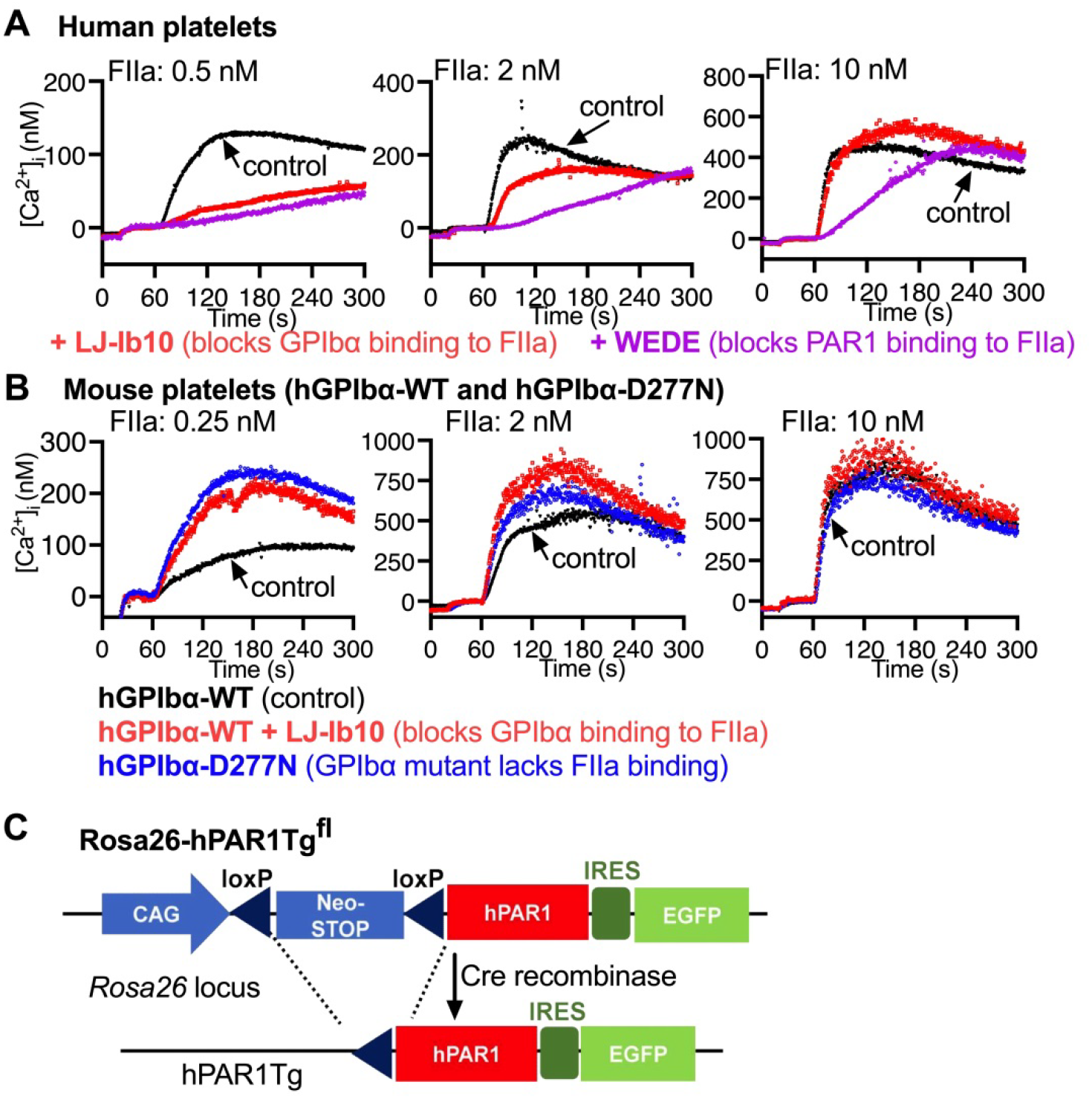
Different effects of FIIa binding to GPIbα on Ca^2+^ signaling in human and mouse platelets. Washed platelets were prepared from human healthy donors (**A**) or mice expressing on the platelet surface hGPIbα-WT or hGPIbα-D277N defective in FIIa binding (**B**). These platelets were loaded with 5 μM Fura 2-AM to measure changes in intracytoplasmic Ca^2+^ concentration after stimulation with different concentrations of FIIa in the absence or presence of MoAb LJ-Ib10, which binds to human GPIbα interfering with FIIa-dependent platelet activation. WEDE, blocking PAR1 function, was used only with human platelets. (**C**) Schematic representation of the targeting construct used for creating a mouse strain expressing hPAR1.

### Expression of hPAR1 on mouse megakaryocytes (MKs) and platelets

In this study, unlike in previous attempts^26,27^, we first generated Rosa26-hPAR1Tg^fl^ mice that were then bred with PF4-Cre transgenic mice to excise the floxed STOP cassette in the megakaryocyte (MK) lineage,^28^ resulting in mice whose platelets expressed hPAR1Tg along with EGFP (Figure 2A). To assess the functionality of expressed hPAR1, washed platelets prepared from hPAR1Tg mice or human healthy donors were stimulated by the specific PAR1 activating peptide, TFLLRN-NH_2_ (P1-AP). At 10 μM concentration, this induced similar aggregation in platelets from hPAR1Tg mice and human controls (Figure 2B). To avoid possible confounding effects by mouse PAR3, we crossed hPAR1Tg (hP1Tg) and PAR3^−/−^ mice (P3^−/−^) to obtain the P3^−/−^hP1Tg strain, whose platelets expressed ∼30-40% hPAR1 as compared to human platelets, but ∼80% after correction for the larger size of the latter (Figure 2C-D). Western blotting of platelet lysates confirmed the expression of hPAR1 in P3^−/−^hP1Tg platelets, likely glycosylated judging from the apparent molecular mass (Figure 2E, red arrows). We also examined hPAR1 expression^34^ in P3^−/−^hP1Tg BM MKs and found very low levels up to 8N cells, increasing to a maximum in 64N cells; expression of integrin αIIb (CD41), evaluated as a control, followed a similar pattern (Figure 2F).

**Figure 2.**
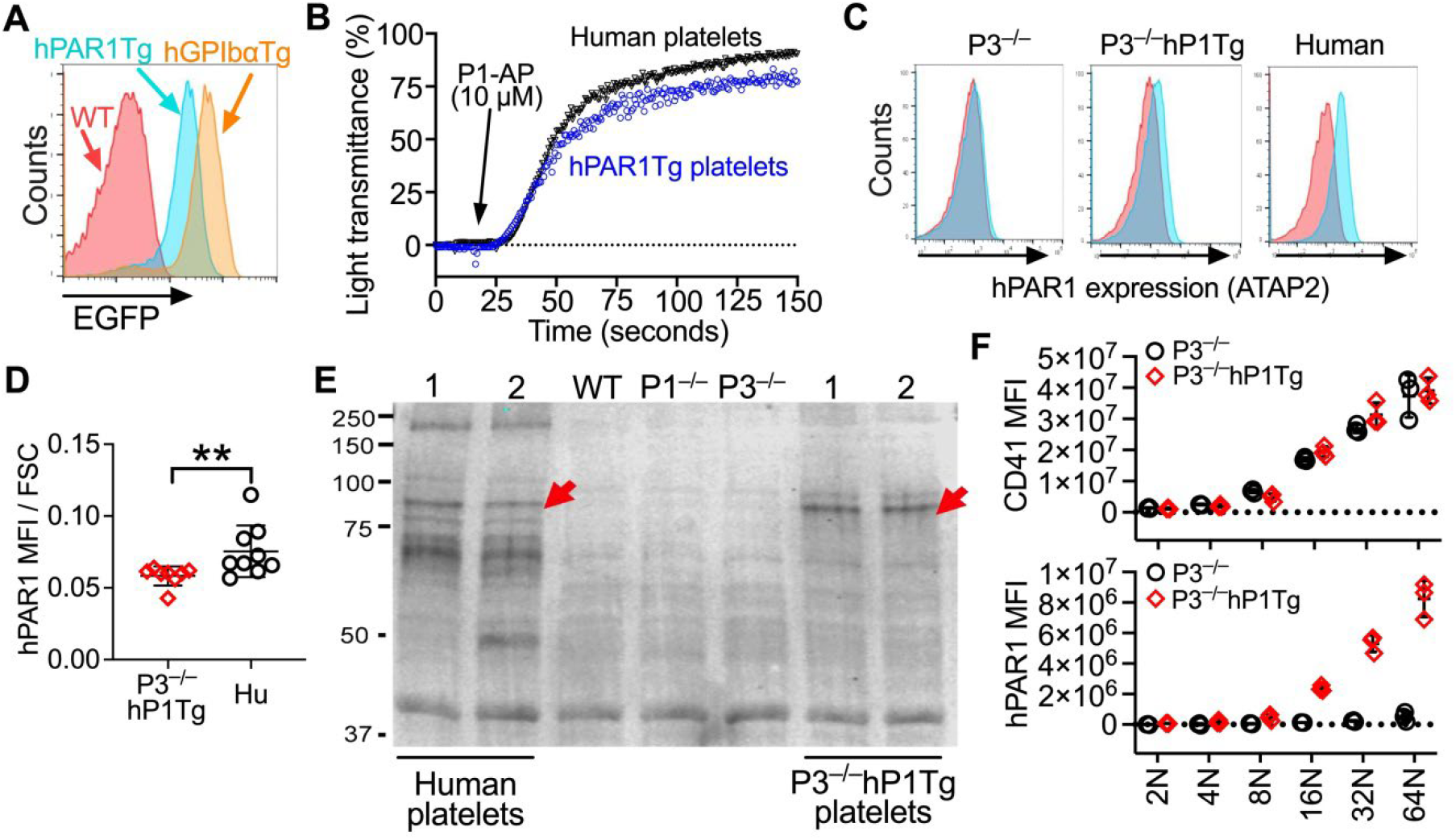
Expression of hPAR1 on mouse platelets. (**A**) Identification by flow cytometry of mouse platelets expressing EGFP and hPAR1Tg or hGPIbαTg generated using the same targeting vector. (**B**) Aggregation of hPAR1Tg mouse and human washed platelets induced by 10 μM P1-AP. (**C**) Expression of hPAR1 in P3^−/−^hP1Tg compared to human platelets analyzed by flow cytometry using the anti-hPAR1 antibody ATAP2. (**D**) Expression of hPAR1 on platelets from human controls and P3^−/−^hP1Tg mice measured as mean fluorescence intensity (MFI) of bound antibody divided by forward side scatter (FSC) to correct for the difference in platelet size. (**E**) Western blotting of platelet lysates after reaction with the anti-hPAR1 antibody ATAP2; red arrows point to the hPAR1 signal in human and P3^−/−^hP1Tg platelets. (**F**) BM cells were prepared from P3^−/−^ and P3^−/−^hP1Tg mice (n = 3 in each group) to analyze by flow cytometry surface expression of integrin αIIb (CD41) and hPAR1 on MKs of different ploidy. Results are shown as scatter dot plots with mean ± SD.

### Aggregation of P3^−/−^hP1Tg mouse platelets in response to P1-AP

We reassessed the functionality of hPAR1 on P3^−/−^hP1Tg mouse platelets by measuring aggregation of washed platelets (3·10^5^/μL) in response to P1-AP stimulation and found a dose-dependent response (Figure 3A) similar to that of human control platelets (Figure 3B). The P1-AP concentration required to induce full aggregation was ∼2.5-5 μM for human as compared to 7.5 μM for P3^−/−^hP1Tg platelets, compatible with lower receptor number or function. Importantly, maximal aggregation of P3^−/−^hP1Tg platelets induced by 7.5 μM P1-AP was inhibited by the human PAR1 antagonist drug, vorapaxar, ^38^ dose-dependently in the 40-320 nM concentration range (Figure 3C), compared to 80-160 nM with human platelets activated by 2.5 μM P1-AP (Figure 3D). Consistent with signaling mediated by hPAR1, dose-dependent AKT phosphorylation was evident in P3^−/−^hP1Tg platelets as well as in human platelets (Figure 3E) stimulated by P1-AP and confirmed by quantification of band intensities (Figure 3F, top panel). A similar pattern of dose-dependent phosphorylation was observed with ERK, but the change after stimulation relative to background was smaller (Figure 3F, bottom panel).

**Figure 3.**
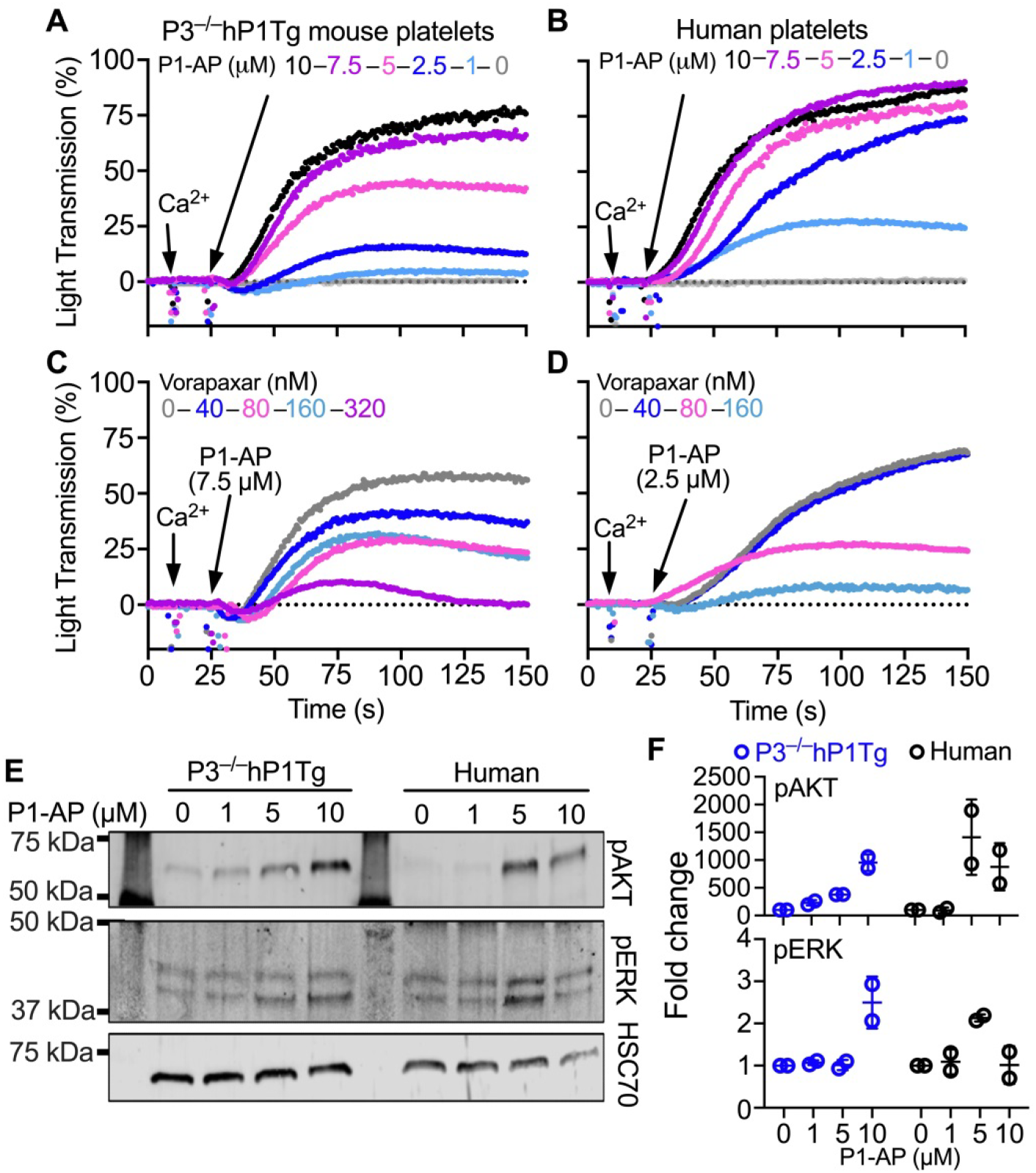
Functional analyses of hPAR1 in P3^−/−^hP1Tg mouse platelets stimulated by PAR1 activation peptide (P1-AP). Aggregation induced by different concentrations of P1-AP was evaluated in washed platelet suspensions prepared from P3^−/−^hP1Tg mice (**A**) or healthy human donors (**B**). The inhibitory effect of the PAR1 antagonist drug, vorapaxar, was examined using the minimum P1-AP concentration enabling full aggregation, corresponding to (**C**) 7.5 μM for P3^−/−^hP1Tg platelets and (**D**) 2.5 μM (for human platelets. (**E**) Washed platelets prepared from P3^−/−^hP1Tg mice or human healthy donors were stimulated with different concentrations of P1-AP and platelet lysates were analyzed by Western blotting. HSC70 was used as a loading control. (**F**) The signal intensity of protein bands as in (B) was quantified with the Licor ImageStudio instrument. These experiments were repeated two times and the quantified results (mean ± SD) are shown as scatter plots with mean value.

### Characterization of P3^−/−^hP1Tg mouse platelets response to FIIa stimulation

Compared to P3^−/−^ platelets expressing PAR4 as the sole FIIa receptor (Figure 4A), P3^−/−^hP1Tg platelets expressing hPAR1 in addition to PAR4 (Figure 4B) exhibited a trend toward enhanced response to FIIa at low (0.6-0.8 nM) concentration, although the difference did not reach statistical significance (Figure 4C). Similar aggregation response was observed with P3^−/−^ and P3^−/−^hP1Tg platelets at higher FIIa (1.2-2.4 nM). To ascertain further the relative contribution of hPAR1 and mouse PAR4 to FIIa-induced activation of P3^−/−^hP1Tg platelets, we used the PAR4-specific inhibitor, BMS-986120 (BMS)^41^. At 80 nM concentration, BMS completely inhibited the aggregation of P3^−/−^ platelets up to 1.6 nM FIIa (Figure 4D). In contrast, with the same BMS concentration, P3^−/−^hP1Tg platelets exhibited residual aggregation, albeit reduced, in response to 1.6 and 1.2 nM FIIa (Figure 4E). This residual aggregation in the presence of BMS-986120 was completely abolished by the PAR1 inhibitor vorapaxar (Figure 4F), consistent with response to FIIa stimulation after PAR4 blockade resulting from hPAR1-mediated signaling on mouse platelets.

**Figure 4.**
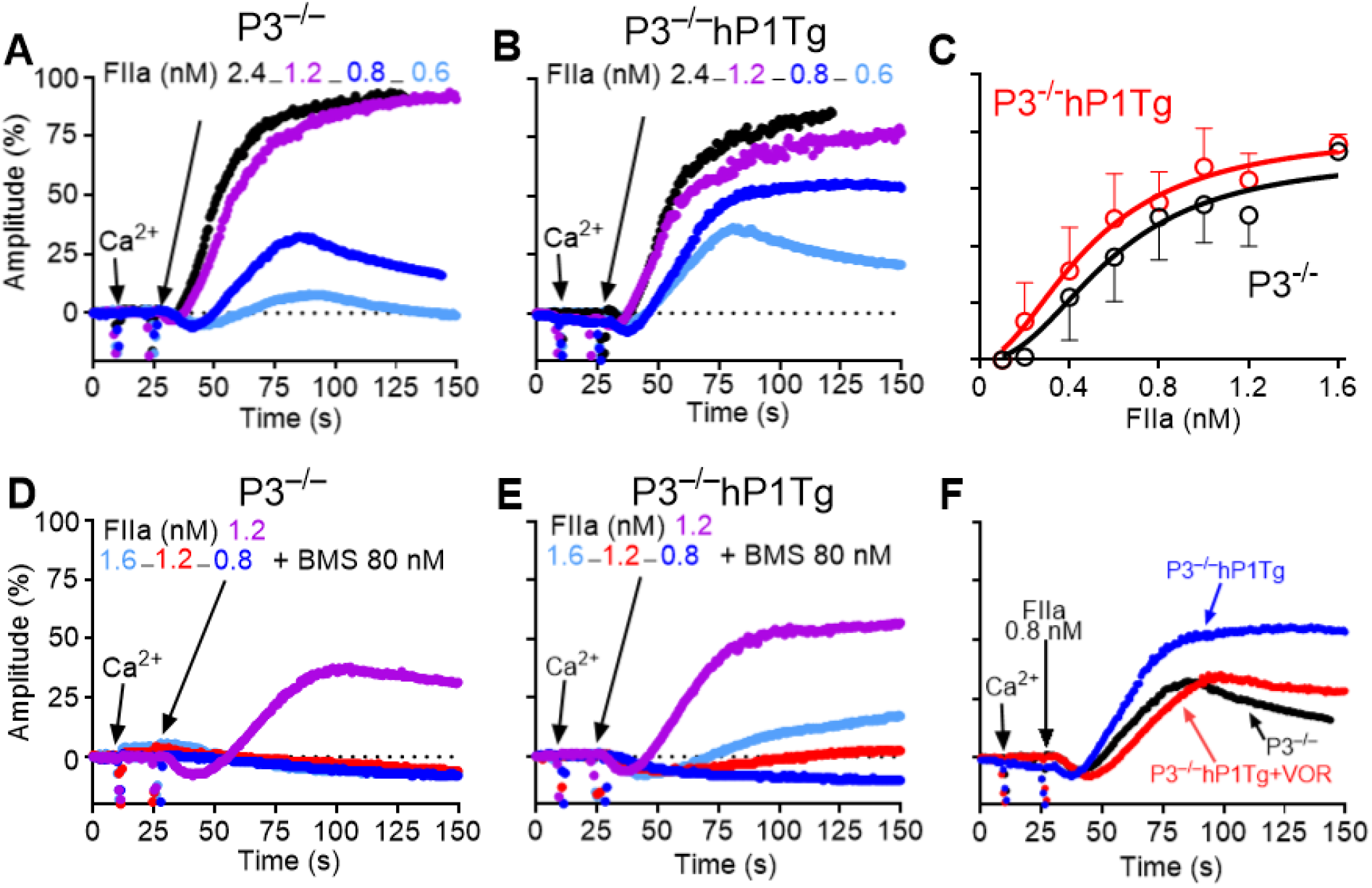
Aggregation of P3^−/−^hP1Tg as compared to P3^−/−^ mouse platelets stimulated by α-thrombin (FIIa). Washed platelets from P3^−/−^ (**A**) and P3^−/−^hP1Tg (**B**) mice were induced to aggregate by various FIIa concentrations. Mean ± S.E.M. of amplitude from six independent biological experiments at the indicated FIIa concentration (**C**). (**D, E**) Aggregation of P3^−/−^ and P3^−/−^hP1Tg mouse platelets were induced by FIIa in the presence of the PAR4 inhibitor, BMS-986120 (BMS, 80 nM). Note the complete aggregation inhibition by BMS of P3^−/−^ platelet, but only partial of P3^−/−^hP1Tg platelets at the two higher FIIa concentrations. (**F**) The residual aggregation of P3^−/−^hP1Tg compared to P3^−/−^ platelets induced by 1.6 nM FIIa in the presence of BMS was abolished by vorapaxar (VOR; 400 nM).

### PAR1 and PAR4 response to FIIa stimulation in human and P3^−/−^hP1Tg platelets

We used vorapaxar^39,40^ and BMS-986120 to compare PAR1 and PAR4 function in human and P3^−/−^hP1Tg mouse platelets stimulated with a fixed (1.6 nM) FIIa concentration. In human platelets, BMS-986120 alone in the final concentration range of 80 nM to 8 μM had minimal effect on FIIa-induced aggregation, as did vorapaxar at 400 nM; however, combination of 400 nM vorapaxar with 80 nM BMS-986120 caused profound aggregation inhibition (Figure 5A). In contrast, in P3^−/−^hP1Tg mouse platelets, BMS-986120 dose-dependently inhibited aggregation, which was abrogated at the concentration of 160 nM. Vorapaxar at 400 nM had no effect by itself but added it to the partially inhibitory dose of 80 nM BMS-986120 completely abrogated aggregation, with effect comparable to 160 nM BMS-986120 alone (Figure 5B).

**Figure 5.**
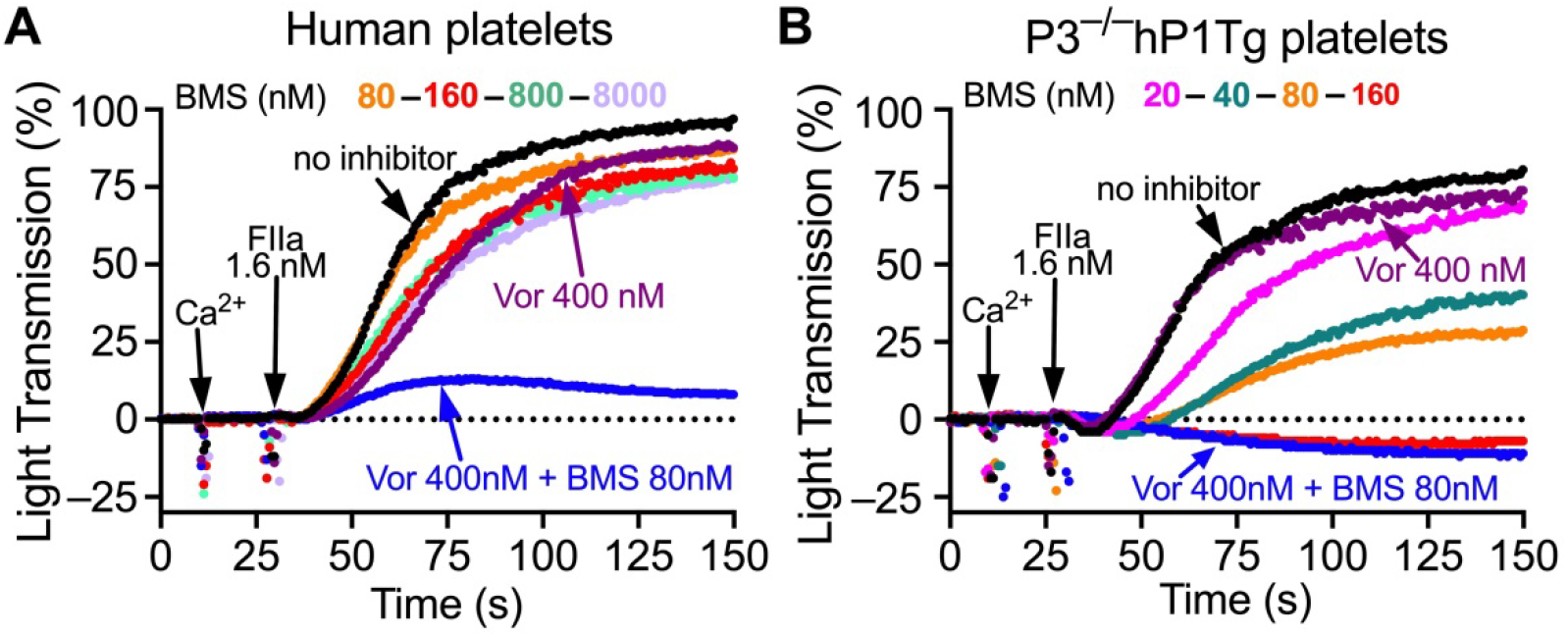
Effect of PAR1 and PAR4 inhibition on FIIa-induced aggregation of human and P3^−/−^hP1Tg mouse platelets. Washed platelets prepared from healthy donors (**A**) and P3^−/−^hP1Tg mice (**B**) were stimulated with 1.6 nM FIIa in the presence of different concentrations of BMS-986120 (BMS), vorapaxar (VOR) or both and evaluated for aggregation.

### Bleeding time of hPAR1 expressed in P3^−/−^hP1Tg mice

As previously reported,^29^ P3^−/−^ mice present a defect of hemostasis reflected in variably prolonged bleeding after clipping of the tail extremity. This defect was completely rescued by hPAR1 expression in P3^−/−^hP1Tg mice and rescuing was abrogated by pre-treating the latter with vorapaxar inhibiting PAR1 function (Figure 6).

**Figure 6.**
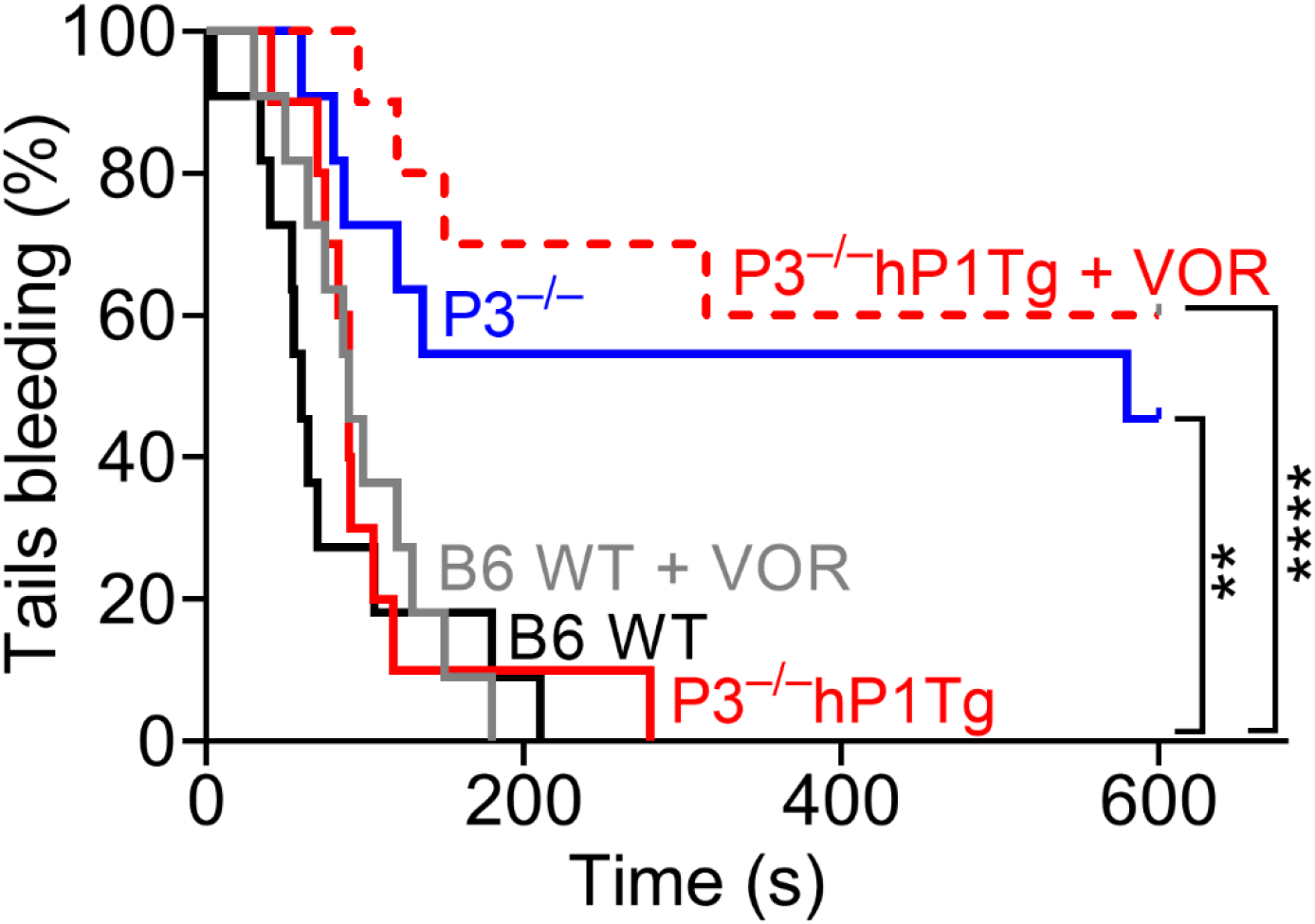
Hemostasis after tail clipping in P3^−/−^hP1Tg and control mice. (**A**) The arrest of bleeding after tail clipping was evaluated in C57BL/6J wild type (B6 WT), P3^−/−^ and P3^−/−^hP1Tg mice; in B6 WT and P3^−/−^hP1Tg mice the test was also performed 5 minutes after intravenous administration of vorapaxar (3.6 mg/kg). N = 10 in each group. The percentage of tails bleeding within 600 seconds after injury is shown by Kaplan-Meier analysis, with differences between groups evaluated by the log-rank (Mantel-Cox) test with Bonferroni correction for pairwise comparison. ^*^P<0.01. P = 0.012 for the other bracketed comparison.

### The effect of hPAR1 expression on vascular occlusion

To evaluate functional contribution of hPAR1 expressed on mouse platelets to arterial thrombus formation, mice were tested by FeCl_3_ artery injury model.^30^ While only 50% of the mice lacking Par3 (P3^-/-^) were able to generate a thrombus able to firmly occlude the carotid artery, five out of six mice expressing hPAR1 (P3^−/−^hP1Tg) did so; pharmacological treatment with Vorapaxar inhibited stable occlusion only in mice expressing hPAR1 but not in B6 mice (Figure 7).

**Figure 7.**
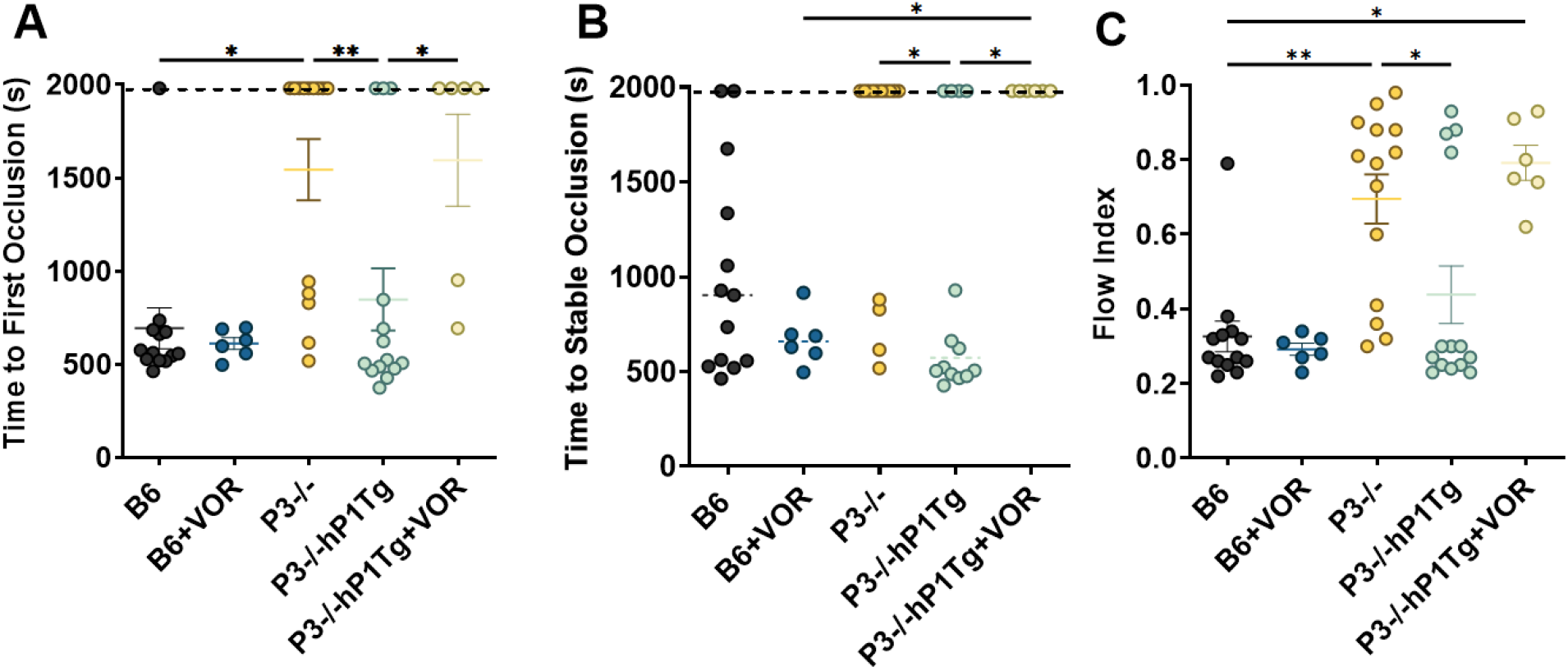
FeCl_3_-induced carotid artery occlusion model. Time to first occlusion (A) and time to stable occlusion (B) were measured following ferric chloride (0.22 M) induced injury to the left carotid artery. Flow index (C) was calculated as described in Materials and Methods. Indicated groups were treated with vehicle or vorapaxar (VOR) (3.6 mg/kg). Censored values (1980 s) indicate absence of occlusion within the recording window (33’). Statistical differences were determined using Kruskal-Wallis test with Dunn’s multiple comparisons. ^*^P < 0.05, ^**^P < 0.01.

## Discussion

Platelet activation by thrombin (FIIa) mediated by PARs is a key event in both host defense and pathogenic mechanisms^1,5,31^; and is fundamentally different in humans and mice. Human platelets express PAR1 and PAR4 that independently can generate signals activating platelets; mouse platelets express PAR3 and PAR4, with PAR3 acting as a co-receptor that facilitates PAR4 cleavage by FIIa but lacks independent signaling function^1,31,32^. This fundamental difference has made it difficult to translate directly results from murine models to human platelet physiology.^26,27,37^

The results of this study demonstrate the generation of a transgenic mouse, designated P3−/−hP1Tg, whose platelets express human PAR1 along with endogenous mouse PAR4 in the absence of murine PAR3^27^, allowing us to evaluate the role of human PAR1 in thrombin-induced platelet activation, hemostasis, and vascular occlusion^29^. Unlike previous attempts that failed to generate functional hPAR1 expression in murine platelets^26^, this strategy successfully achieved stable and platelet-specific hPAR1 expression by targeted integration into the Rosa26 locus, a well-established safe harbor site for transgene expression^23^. This allowed us to evaluate not only whether hPAR1 could functionally compensate for the absence of PAR3 but also whether its presence could support thrombus formation in vivo and respond appropriately to pharmacological inhibitors. A major finding of this study is that hPAR1 is sufficient to restore hemostasis and thrombosis in P3−/− mice^29^. In tail bleeding time assays, which serve as a surrogate for primary hemostasis, hPAR1 expression fully corrected the prolonged bleeding time observed in P3−/− mice, indicating that human PAR1 can functionally replace murine PAR3 in thrombin-dependent platelet activation^27^. Furthermore, in the FeCl_3_-induced arterial occlusion model, P3−/−hP1Tg mice exhibited stable occlusive thrombus formation at rates comparable to WT mice. Importantly, thrombosis was inhibited by vorapaxar^42^, confirming that thrombus formation in this model was dependent on PAR1 activation. These results establish P3−/−hP1Tg mice as a functionally validated, pharmacologically responsive in vivo system for studying human PAR1 in thrombin-dependent platelet activation. A surprising aspect of our findings was that thrombin-induced platelet aggregation was minimally different between P3−/− and P3−/−hP1Tg mice, despite the clear impact of hPAR1 on hemostasis and thrombosis^36^. This observation is in contrast with expectations based on human platelet physiology, where PAR1 is the primary thrombin receptor^3,5^ and mediates rapid aggregation responses. One possible explanation for this discrepancy is that in mouse platelets, PAR4 plays a dominant role in thrombin-induced aggregation, such that the presence or absence of PAR1 has a limited impact on ex vivo aggregation assays. In contrast, the ability of hPAR1 to restore hemostasis and support thrombus formation in vivo suggests that its role may extend beyond initial aggregation, possibly by enhancing thrombus stability, platelet procoagulant activity, ^46^ or platelet-fibrin interactions. Further studies will be required to determine whether hPAR1 alters platelet procoagulant responses, fibrin deposition, or clot retraction, any of which could explain why hPAR1 restores in vivo thrombosis despite a limited effect on aggregation. Additionally, it remains to be determined whether hPAR1 influences secondary signaling pathways involved in sustained platelet activation, such as PAR1-PAR4 crosstalk, which has been proposed in human platelets.^35^ Given that murine PAR4 and human PAR4 exhibit distinct regulatory features, the extent to which hPAR1 modifies thrombin signaling in a mouse platelet environment warrants further investigation. Some of the key remaining questions are the nature and the consequences of the functional interaction between hPAR1 and murine PAR4 in P3−/−hP1Tg platelets. In human platelets, PAR1 and PAR4 function independently^3,43^, with distinct activation tempos: due to effective GPIbα-mediated funneling, minimal thrombin is sufficient to cleave PAR1, triggering a rapid but transient response.^13^ In contrast, PAR4 requires higher thrombin concentrations for full activation and delivers sustained signaling. This kinetic difference is believed to contribute to quick and stable platelet activation ensuring a prolonged hemostatic response. However, in mouse platelets, PAR3 is required to facilitate thrombin cleavage of PAR4, which does not have a high affinity exosite I binding site for thrombin. In P3−/−hP1Tg platelets, hPAR1 is now the putative high-affinity thrombin receptor, effectively bypassing the need for PAR3 in the thrombin activation cascade. However, since PAR4 remains of murine origin, it is unclear whether it functions in the same manner as human PAR4 in sustaining platelet activation. Our data suggest that hPAR1 may not fully recapitulate its independent role seen in human platelets but instead supports PAR4 activation in a manner that sustains in vivo hemostasis and thrombosis. Previous studies have suggested that PAR1 and PAR4 may form heterodimers in human platelets, leading to altered cleavage dynamics and intracellular signaling cascades. If such interactions occur between hPAR1 and murine PAR4 in our model, they could explain why hPAR1 restores in vivo thrombosis without significantly altering aggregation. Additionally, it has been proposed that mouse PAR4 exhibits differences in its C-terminal tail that affect its interaction with intracellular signaling effectors, such as β-arrestin and G-protein subtypes. Further characterization of PAR1-PAR4 interactions in P3−/−hP1Tg platelets will help clarify whether hPAR1 in mice engages with PAR4 in a manner that differs from its function in human platelets. The ability of vorapaxar, an FDA-approved PAR1 inhibitor, to effectively block thrombosis in P3−/−hP1Tg mice is a critical validation of the model’s relevance. Our results now demonstrate that P3−/−hP1Tg mice provide a functional and pharmacologically relevant system for assessing PAR1-directed antithrombotic agents. This model also allows for the evaluation of next-generation PAR1 inhibitors with improved safety profiles. While vorapaxar has shown efficacy in preventing thrombotic events in patients with cardiovascular disease, its use is limited by an increased risk of bleeding, particularly in patients with a history of stroke.^44^ Developing new PAR1 inhibitors that selectively target platelet activation while preserving endothelial PAR1 function has been a major goal in drug development. P3−/−hP1Tg mice could serve as a valuable vivo screening platform for testing novel PAR1 inhibitors with enhanced specificity and reduced bleeding risk. PAR1 is not only a key mediator of platelet activation but also plays significant roles in vascular biology, inflammation, and coagulation-dependent immune responses.^45^ The P3−/−hP1Tg model provides a unique tool for testing how hPAR1 modulates platelet interactions with immune cells, endothelial activation, and inflammatory cytokine release in vivo. In summary, we have developed and characterized a novel transgenic mouse model expressing human PAR1 in platelets, providing a unique preclinical platform for studying thrombin signaling and evaluating PAR1-targeting therapies.

## Acknowledgements

The authors thank Laurent Mosnier at the Scripps Research Institute for providing human PAR1 cDNA and Greg Martin and Sergey Kupriyanov at Mouse Genetics Core Facility of the Scripps Research Institute for generation of the transgenic mice. This work was supported by National Institutes of Health, National Heart, Lung, and Blood Institute grants HL-135294 (ZMR), HL-129011 (TK), by fellowships and additional financial support from MERU Foundation (Italy) to TK, SK, AZ, YM, and RA; and by the National Foundation for Cancer Research (SK and TK).

## Authorship Contributions

S.K. generated hPAR1Tg mice, performed experiments and co-wrote the manuscript. A.Z. and R.A. performed platelet aggregation and Ca^2+^ assays and contributed to data analysis and interpretation Y.M. performed flow cytometry analysis. A.Z. performed *in vivo* animal study. E.W. performed Western blotting and data analysis. Z.M.R supervised the project, helped with experimental design and data analyses and co-wrote the manuscript. T.K. co-designed the study, performed experiments, analyzed data and co-wrote the manuscript.

